# (3*S*,6*E*)-Nerolidol-mediated rendezvous of Cyclocephalini beetles, *Cyclocephala paraguayensis*, in bottle gourd flowers

**DOI:** 10.1101/2020.06.09.142059

**Authors:** Arodí P. Favaris, Amanda C. Túler, Weliton D. Silva, Sérgio R. Rodrigues, Walter S. Leal, José M. S. Bento

## Abstract

Cyclocephalini beetles of genus *Cyclocephala* (Coleoptera: Melolonthidae: Dynastinae) use flowers of some plants as food, shelter, and mating stands. However, little is known about floral scent chemistry involved in this interaction. Here we show that a sesquiterpene alcohol mediates attraction of *Cyclocephala paraguayensis* Arrow on bottle gourd flowers, *Lagenaria siceraria* (Curcubitaceae). Both males and females started to aggregate on flowers at twilight; after that, mating began and remained for the entire night. The major constituent of the airborne volatiles from *L. siceraria* was fully characterized as (3*S*,6*E)*-nerolidol, which elicited electroantennographic responses on male and female antennae. Field bioassays showed that traps baited with the natural stereoisomer or a mix of nerolidol isomers captured significantly more males and females of *C. paraguayensis* than control traps. Analysis from the gut content of these Cyclocephalini beetles showed the presence of pollen, suggesting that they also use bottle gourd flowers for their nourishment. Taken together, these results suggest that (3*S*,6*E)*-nerolidol plays an essential role in the reproductive behavior of *C. paraguayensis* by eliciting aggregation, mating, and feeding.

## Introduction

Floral scents have been reported as important assets for the visitation of beetles of the tribe Cyclocephalini (Melolonthidae: Dynastinae) [1–4]. The interaction of beetles and flowers may result in mutualistic pollination typically by the cantharophilous floral syndrome of basal angiosperms, whose flowers are usually bisexual, protogynous, very scented, and thermogenic [4,5]. The Cyclocephalini beetles are primarily attracted by strong odors released by flowers as chemical cues to indicate a rendezvous site for food, shelter, and mating stands [4,6,7].

The floral volatile organic compounds (VOCs) in the plant families of Araceae, Annonaceae, Arecaceae, and Magnoliaceae, have been demonstrated to be comprised of pyrazines, ketones, esters, and methoxystyrenes [8–13]. By contrast, VOCs produced by eudicot flowers that are visited by Cyclocephalini beetles of the genus *Cyclocephala* have never been identified.

*Cyclocephala paraguayensis* Arrow, is a Neotropical species that is frequently found on eudicot plant species [14,15]. Here, we observed this beetle visiting flowers of *Lagenaria siceraria* (Molina) Standle (Curcubitaceae), which incidentally represents a new host flower record for the beetle family Melolonthidae. This plant is originated from Africa and nowadays is spread worldwide [16]; it produces white flowers with separate sexes in the same plant (monoecious) but can assume andromonoecious form, when hermaphrodite flowers are present [17]. Anthesis occurs throughout the night, at which time the lepidopteran pollinators, specially hawkmoths, visit the flowers [18].

We surmised that the volatile components of bottle gourd flowers mediate aggregative behavior of *C. paraguayensis* for feeding and mating. We studied the floral scent chemistry of *L. siceraria* flowers and showed that (3*S*,6*E)*-nerolidol plays an important role in the reproductive behavior of *C. paraguayensis* by eliciting aggregation, mating, and feeding.

## Results

### Sexual and feeding behavior of insects

The behavior of *C. paraguayensis* visitation on *L. siceraria* flowers was observed under field conditions at night for four hours (7:00 p.m to 11:00 p.m.). Beetles activity started by the twilight (∼7:30 p.m.), when males of *C. paraguayensis* were the first to land on *L. siceraria* flowers (n = 36; two-tailed Exact Binomial test; *P* = 0.011). At 8:00 p.m., the number of visiting males and females was 18.50 ± 5.97 (mean ± SD), increasing until 11:00 p.m. to 26.75 ± 7.46. Paired males and females of *C. paraguayensis* (2.25 ± 3.2) were present on bottle gourd flowers from 8:00 p.m. until the end of the observation period. Both sexes of *L. siceraria* flowers were visited. Seventy-five percent of bottle gourd flowers harbored more than one mating couple, thus, suggesting that aggregation on bottle gourd flowers created mating opportunities for the beetles.

These Cyclocephalini beetles usually remained inside the floral receptacles until the next morning (Fig 1A-B). Bottle gourd flowers presented gnawing marks or even major damages resulted from beetles feeding (Fig 1E-F). Of note, pollens of these flowers were found in the gut of beetles (80% females; 50% males, n = 10) when dissected (Fig 1G-H).

**Fig 1.**
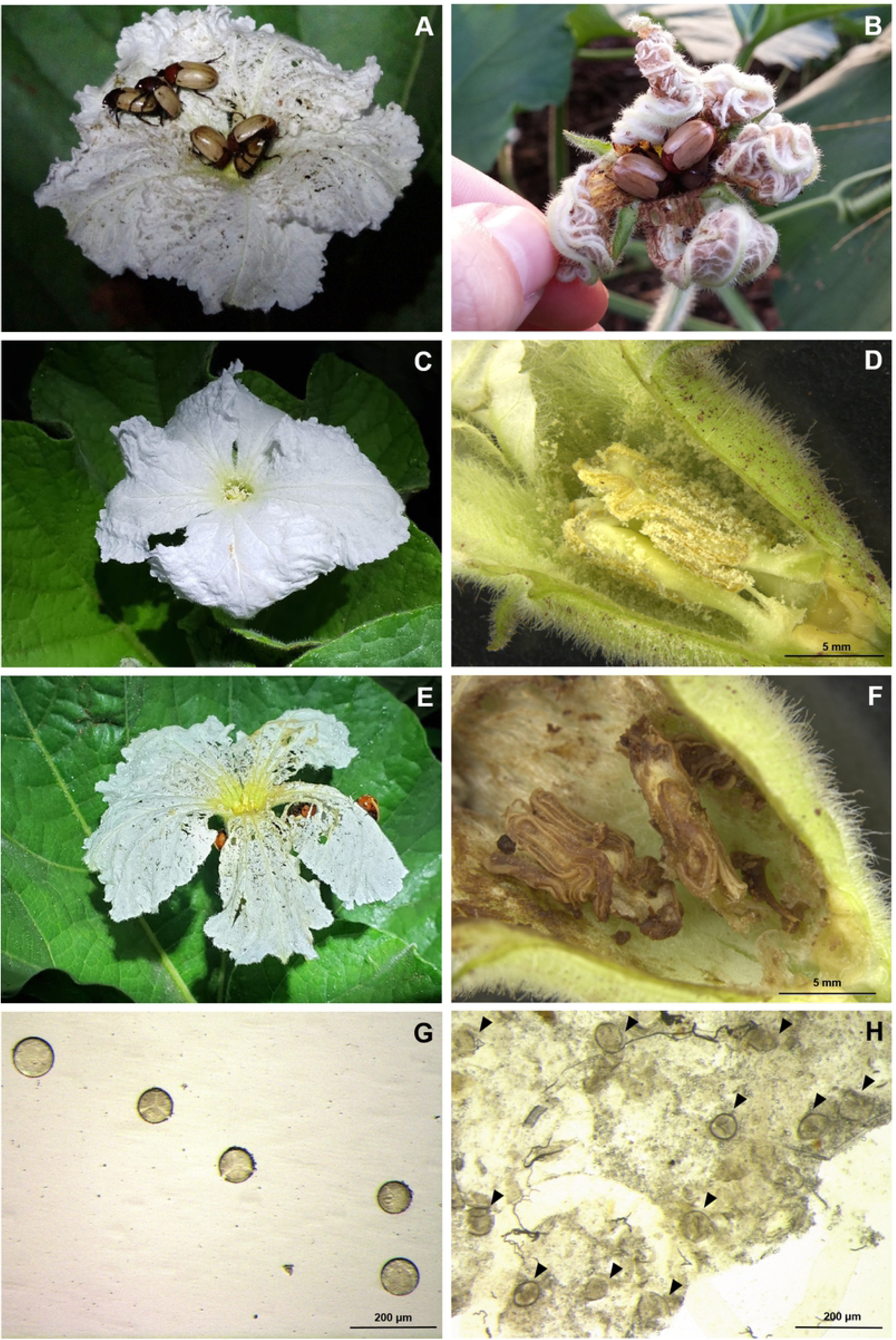
Sexual and feeding behavior of Cyclocephalini beetle, *Cyclocephala paraguayensis*, in bottle gourd flowers, *Lagenaria siceraria*. A) Aggregating and mating on flowers at night; B) *C. paraguayensis* inside senescent flowers in the next morning of beetle’s attraction; C) Non-damaged *L. siceraria* flower in the field; D) Stereoscopic cut view of a non-damaged *L. siceraria* flower; E) *L. siceraria* flower damaged by *C. paraguayensis* in the field; F) Stereoscopic cut view of a damaged flower showing beetle gnawing and excrement signal as a result of feeding activity; G) Fresh pollen grain of *L. siceraria*; H) Pollen grains of *L. siceraria* extracted from *C. paraguayensis* gut (black triangles indicate the pollen grains).

### Floral scent identification

Gas chromatography linked to electroantennographic detection (GC-EAD) analysis of airborne volatiles collected from *L. siceraria* showed the presence of a prominent, antennally active peak at 20.5 min (Fig 2).

**Fig 2.**
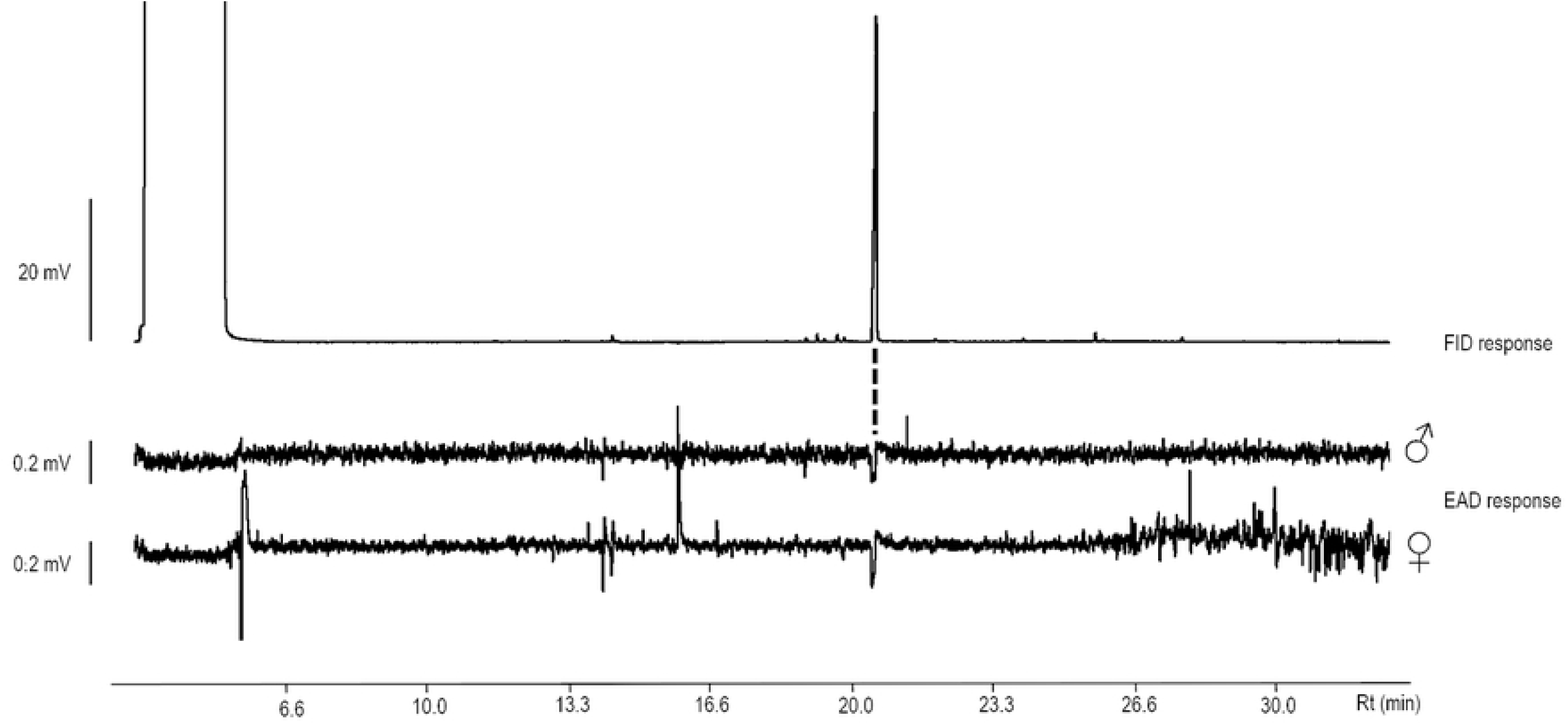
GC-EAD analysis of floral scents extract of *Lagenaria siceraria* by using male and female antenna of Cyclocephalini beetle, *Cyclocephala paraguayensis*, as sensing elements. “FID” flame-ionization detection.

The prominent compound was tentatively identified by GC-MS as a nerolidol isomer, which led us to conduct *cis* and *trans* identification followed by chiral analysis. The EAD active compound was then identified as *trans*nerolidol (i.e., (6*E*)-nerolidol, IR = 1568) by comparing the mass spectra and retention indexes of the natural product and authentic standards (Fig 3a-f). Lastly, chiral analysis using nerolidol standards led to the full characterization of the natural product as (*3S,6E*)-nerolidol (Fig 3g).

**Fig 3.**
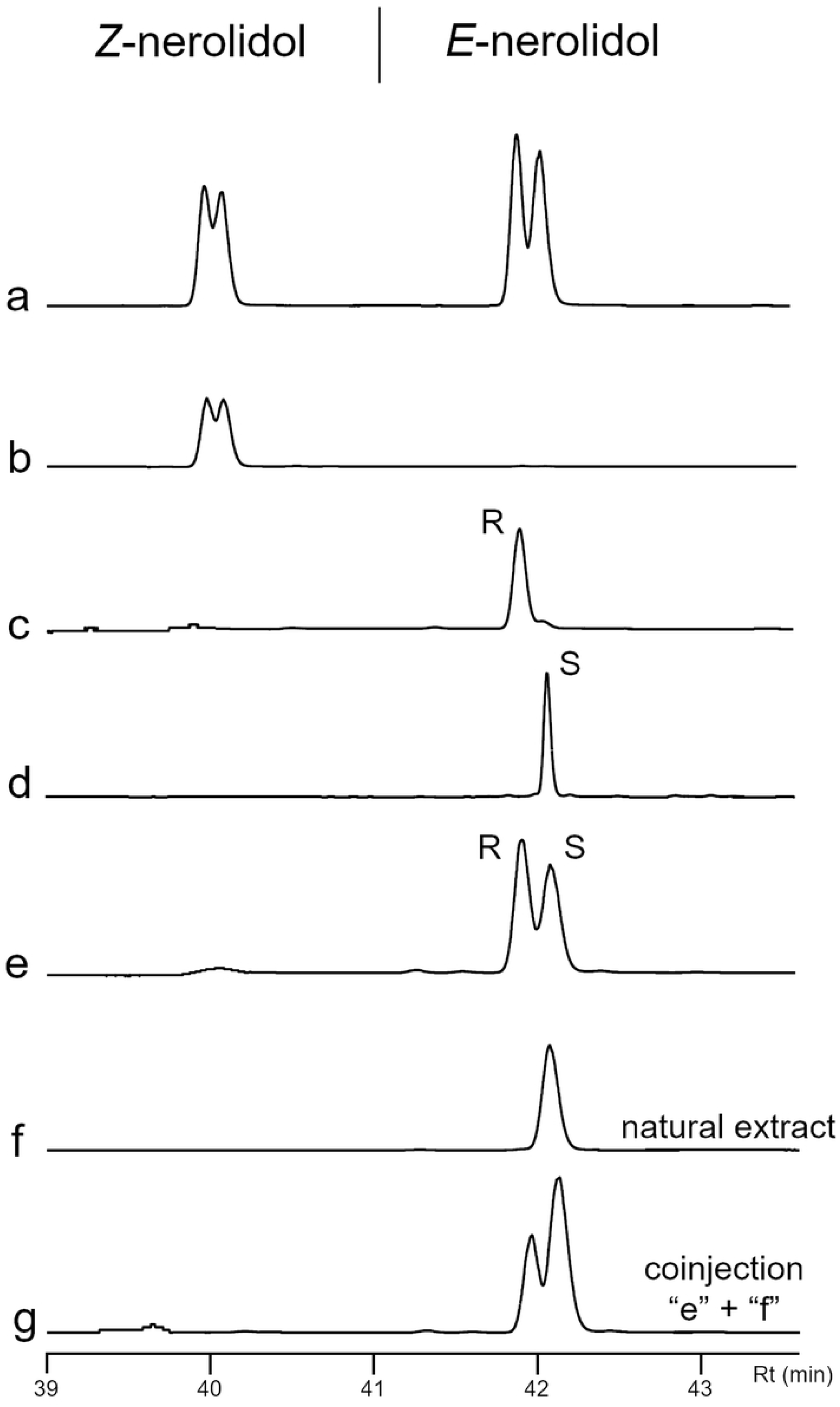
Analysis of bottle gourd flowers, *Lagenaria siceraria*, extract, and synthetic isomers of nerolidol by gas chromatography equipped with a chiral column. a) *Z* and *E*-nerolidol (left to right) isomers of nerolidol (a mix of isomers); b) *Z*-nerolidol isomers; c) Synthetic (*3R,6E*)-nerolidol (gift from Dr. Robert Hanus, Czech Republic); d) (*E*)-nerolidol from Sigma-Aldrich; e) Mix of 40 ng from each (*3R,6E*)-nerolidol and (*3S,6E*)-nerolidol; f) Natural extract from VOCs of *L. siceraria* flowers; g) Coinjection of “e” and “f” chromatogram extracts.

Quantification of VOCs from male and female flowers showed that (*3S,6E*)-nerolidol comprised 97% of the volatile bouquet and was released at the rate of ∼1.3 μg/hour under field conditions.

### Field evaluation of synthetic nerolidol

In our first field trial, a total of six adults of Cyclocephalini beetle, *C. paraguayensis*, landed on nerolidol-treated dummies, whereas no beetle was observed on control dummies (two-tailed Exact Binomial test; *P* = 0.031).

In the second field experiment, we evaluated the response of adults of *C. paraguayensis* to nerolidol-baited traps in two different locations: Cassilândia, State of Mato Grosso do Sul and Piracicaba, State of São Paulo. In both locations, traps baited with nerolidol (a mix of isomers) captured significantly more beetles than control traps. In Cassilândia, 280 adults of *C. paraguayensis* were captured by nerolidol-baited traps during the tests, with no beetles captured in control traps (two-tailed Exact Binomial test; *P* < 0.001; Fig 4A). Although total captures in Piracicaba (n= 32) were lower than in Cassilândia, nerolidol-baited traps captured significantly more beetles than control traps (two-tailed Exact Binomial test; *P* < 0.001). Interestingly, the sex ratio of captured beetles did not differ significantly from the 1:1 ratio in both Cassilândia (51.4% females; 95% Clopper-Pearson exact confidence interval: 0.454-0.574; *P* = 0.632) and Piracicaba (41.4% females; 95% Clopper-Pearson exact confidence interval: 0.235-0.610; *P* = 0.353).

**Fig 4.**
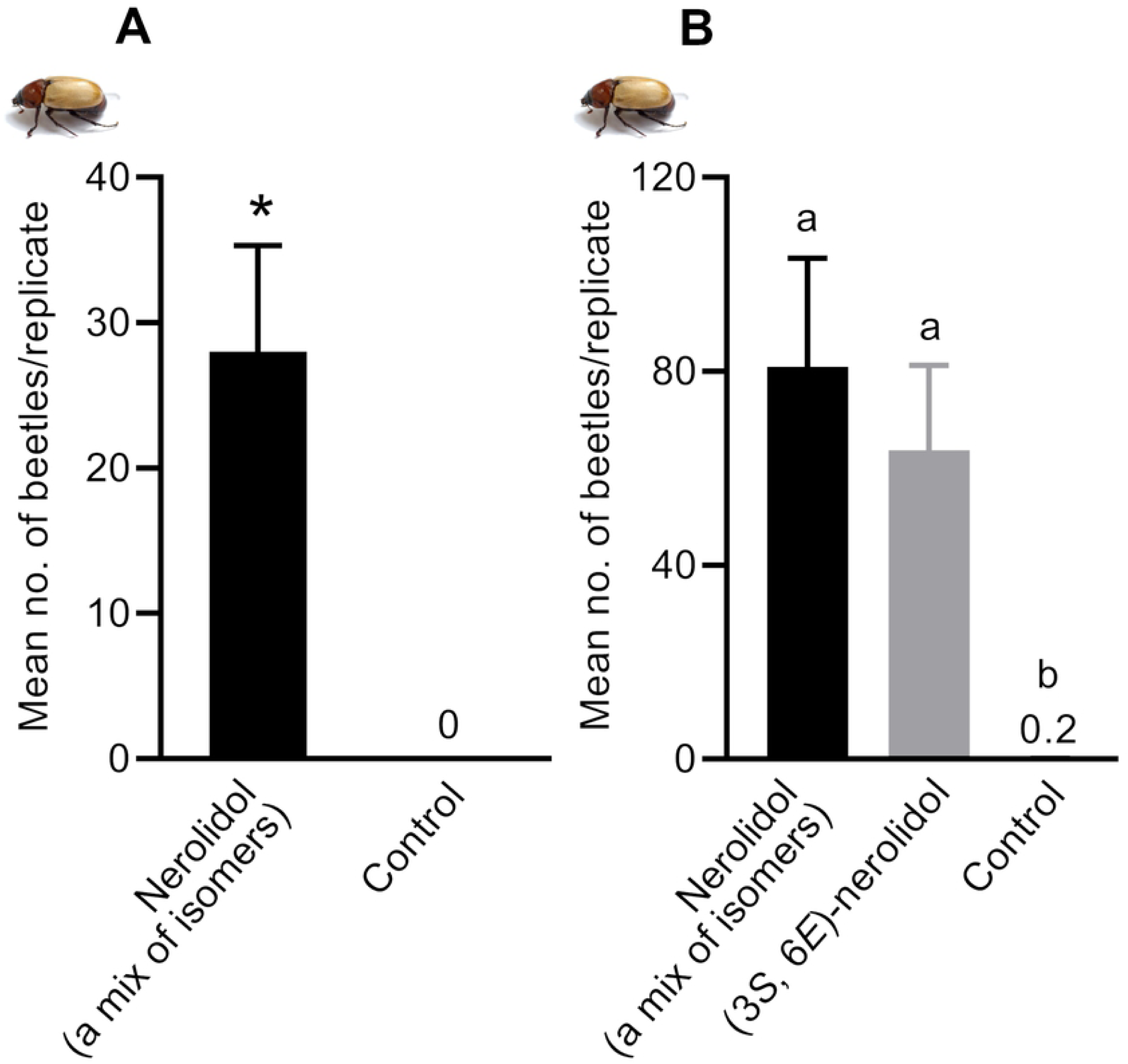
Mean number of Cyclocephalini beetle, *Cyclocephala paraguayensis*, caught per replicate in field experiments conducted in Cassilândia, MS, Brazil. Treatments: Nerolidol (a mix of isomers) = mixture of four isomers of nerolidol; (3*S*, 6*E*)-nerolidol = synthetic nerolidol, which is identical to the stereoisomer of nerolidol produced by *L. siceraria* flowers. A) Mean number of adults of *C. paraguayensis* caught per replicate of nerolidol-baited traps and controls. *Treatment is significantly different at *P* < 0.05 according to Exact binomial test goodness-of-fit; B) Mean number of adults of *C. paraguayensis* caught by traps containing synthetic nerolidol isomers. Treatments had a significant effect on attraction of Cyclocephalini beetles (Fiedman’s Q_2,30_ = 20.002, *P* < 0.001; n = 10 replicates). Means followed by the same letter are not significantly different (REGWQ test, *P* > 0.05).

In the third field tests, we compared captures of *C. paraguayensis* in traps baited with nerolidol (a mix of isomers), (3*S*,6*E*)-nerolidol, and control traps. Throughout these experiments, a total of 1,146 beetles were captured in nerolidol traps (i.e., traps baited with either enantiopure nerolidol, or a mix of isomers). By contrast, only two individuals (males) were caught in control traps. Although treatment and control were significantly different (Q_2,30_ = 20.002; *P* < 0.001; Fig 4B), there was no statistical difference in catches by the mix of isomers or (3*S*,6*E*)-nerolidol (Fig 4B). We, therefore, concluded that the other isomers were neither attractants nor inhibitors. Of note, the sex ratio of the captured beetles in the two nerolidol treatments were somewhat female-biased (mix of isomers = 54.4% female, 95% Clopper-Pearson exact confidence interval: 0.509-0.578; *P* = 0.012; and (3*S*,6*E*)-nerolidol = 58.4% female, 0.544-0.622; *P* < 0.0001). However, as we did not measure the sex ratio of the field populations while conducting bioassays, it is not possible to assert that traps captured significantly more female than male beetles.

## Discussion

Our results suggest that (3*S*,6*E*)-nerolidol mediates aggregation of the Cyclocephalini beetle, *C. paraguayensis*, on bottle gourd flowers, *L. siceraria*. We showed that this sesquiterpene alcohol represents the major VOC emitted by the floral scents of *L. siceraria*, and field tests demonstrated that nerolidol-baited traps captured significantly more beetles than control traps. Surprisingly, field tests showed captures in traps baited with racemic nerolidol (a mix of four isomers) did not differ significantly from those in traps having the enantiopure nerolidol as lure. Thus, the three other non-natural isomers of nerolidol are neither attractant nor deterrent.

We, therefore, concluded that C. *paraguayensis* rendezvous in *L. siceraria* is mediated by nerolidol. There is no evidence in the literature indicating that chemical communication in *C. paraguayensis* involves sex pheromones. If it does, it is unlikely that they need a long-range aggregation-sex pheromone (sensu Cardé 2014) [19] as nerolidol mediates aggregation of both sexes. It is conceivable, however, that *C. paraguayensis* utilizes a hitherto unknown short-range, or contact sex pheromone for species and/or sex recognition.

In this study we showed that *C. paraguayensis* also uses bottle gourd flowers of *L. siceraria* for nourishment given that pollen grains were found in the gut epithelium of males and females. The energy-rich pollen grains may increase the fitness of these Cyclocephalini beetles for mating and oviposition [4]. Our findings do not provide evidence for a mutualistic insect-plant relationship, as previously suggested for other anthophilous cyclocephalines [10,12]. The floral biology traits of curcubits seem to be more suitable for pollination by bees [20], and flowers are particularly successful pollinated by hawkmoths [18]. Additionally, *C. paraguayensis* visit caused damage to *L. siceraria* flowers, although the effects of these damages on the development of fruits remain to be investigated.

## Methods

### Sexual and feeding behavior of insects

The behavior of Cyclocephalini beetle, *C. paraguayensis*, was observed in bottle gourd flowers of *L. siceraria* in a growing area owned by the University of São Paulo, Piracicaba, Brazil (22°42’47.1” S, 47°37’32.7” W). We initially recorded the arrival time of the first beetles on flowers of *L. siceraria*, so the observations in the subsequent days started 30 min earlier. Flowers with insects were marked, and the following parameters were evaluated every 30 min from 7:00 p.m. to 11:00 p.m.: sex of visited flowers; the total number of insects landed per flower; and the number of mating couples per flower. We repeated these observations for 4 d in the same *L. siceraria* area.

Additionally, a sample of 10 males and females of *C. paraguayensis* present within the flowers were taken to verify the presence of pollen in the gut content. Insects were dissected with sharp forceps, and pieces of the midgut and/or the hindgut epithelium were removed and immersed in 0.9% sodium chloride solution. Next, the material was analyzed under a light microscope at 20X magnification. Fresh pollen from *L. siceraria* flowers was prepared in glass slides with the saline solution as reference material.

### Sampling floral VOCs

VOCs from *L. siceraria* flowers were conducted outdoors in the above-mentioned bottle gourd plantation and the laboratory through dynamic headspace extraction. In the field, VOCs were collected from individual male and female *L. siceraria* flowers recognized according to morphological features [17]. These flowers were enclosed within a 17 × 20 cm polyester oven bags (Wyda^®^, Sorocaba, São Paulo, Brazil). Collectors consisted of glass pipets packed with 30 mg of the adsorbent polymer Hayesep^®^ Q (80/100 mesh; Supelco, Bellefonte, PA, USA), which were attached to the bag outlets. Charcoal-filtered air was pushed with a portable pump through the bag and collectors at ∼200 mL/min. Bottle gourd flowers were aerated from 7:30 p.m. to 08:30 p.m., which corresponded to the active period of the beetles. Trapped VOCs were eluted from collectors using 0.3 mL of distilled hexane into 2 mL amber vials that were stored at -30 °C until analysis. Aerations of empty bags were made in parallel to monitor system contaminants. In the same fashion, VOCs were collected from four detached flowers inside a vertical glass jar (55 cm long × 8 cm id) under laboratory conditions (23 ±2 °C; 60 ±10% RH). Collectors attached to the jar outlets were packed with 150 mg of adsorbent polymer. Charcoal-filtered air was pushed through jars and collectors at ∼300 mL/min, and collections were made overnight for up 11 h. Trapped VOCs were eluted from collectors using 1.5 mL of distilled hexane into 2 mL amber vial. To use these extracts for GC-EAD analysis, the resulting aliquots were then concentrated to 500 μL under a gentle flow of N_2_.

### GC- EAD recordings

Extracts of VOCs from *L. siceraria* flowers were analyzed by GC-EAD, i.e., gas chromatography (Shimadzu GC-2010 gas chromatograph, Shimadzu Corp., Kyoto, Japan) linked to an electroantennographic detector system (EAD, Ockenfels Syntech, Buchenbach, Germany) using *C. paraguayensis* antenna as the sensing element. The GC was equipped with a low polarity stationary phase column (30 m × 25 μm × 25 mm; Rtx-5; RESTEK, Bellefonte, PA, USA) connected to a stainless steel crosshead split with two deactivated columns for conducting effluent to FID and EAD detectors at the ratio of 1:3. Helium was used as the carrier gas at a linear velocity of 30.3 cm/s, and 98.3 kPa was set for column head pressure. EAD preparations consisted of male or female excised antennae, and electrodes were made of glass pipette/gold wire filled with a Beadle-Ephrussi Ringer solution (3.75 g NaCl, 0.175 g KCl, 0.14g CaCl_2_·2H_2_O, 500 mL H_2_O). The base of the antenna was inserted into the indifferent electrode, and the tip of antennal lamellae was connected to the recording electrode for signal acquisition (Universal Probe, signal amplification 10x). The antenna preparation was inserted into a glass duct connected to the EAD deactivated column tip, carrying a humidified air stream at 0.3 mL/min. Two microliter aliquots of laboratory-collected volatile extracts were injected in splitless mode at 250 °C, and the GC oven was set at 40 °C for 5 min and then increased to 250 °C at 10 °C/min. FID and EAD signals were recorded by Syntech GcEad software (version 4.6).

### Chemical analysis

The EAD active compound in the VOCs was identified by gas chromatography coupled to a mass spectrometer (GC-MS) with a Shimadzu GCQP-2010 Ultra gas chromatograph (Shimadzu Corp., Kyoto, Japan), equipped with a non-polar stationary phase column (30 m × 25μm × 25 mm; Rxi-1MS; RESTEK, Bellefonte, PA, USA). One microliter of the extract was injected at 250 °C in split mode (1:3). The GC oven temperature was the same for GC-FID/EAD analysis (above). The helium carrier gas was maintained at a flow of 1.3 mL/min in linear velocity at 41.1 cm/s and 70.7 kPa pressure. Both ion source and quadrupole temperatures were set at 250 °C. Mass spectra were recorded in electron impact mode (70 eV) from *m/z* 35-280. Compounds were identified by their retention indexes, mass spectra comparison with Library NIST 11, and injection of authentic standards (Sigma-Aldrich). The target compound was quantified by GC-FID with a Shimadzu GC-2010 gas chromatograph (Shimadzu Corp., Kyoto, Japan) equipped with a non-polar stationary phase column (30 m × 25μm × 25 mm; Rtx-1; RESTEK, Bellefonte, PA, USA) following the same set-up of GC-FID/EAD analysis. We added 3 μL of a 1 mg/mL solution of octadecane as an internal standard in all samples for quantification.

The absolute configuration of flower-produced nerolidol was determined using an Agilent 6890GC with flame ionization detector equipped with a permethylated *β*-cyclodextrin stationary phase chiral column (30 m × 25μm × 25 mm; HP-CHIRAL-20B; J&W Scientific, Folsom, CA, USA). Synthetic nerolidol (a mixture of isomers) and individual geometric isomers were purchased from Sigma-Aldrich to obtain retention time of *cis* and *trans*-nerolidol. A sample of synthetic (*3R,6E*)-nerolidol was kindly donated by Dr. Robert Hanus (Institute of Organic Chemistry and Biochemistry Academy of Sciences of the Czech Republic). All compounds solution in hexane (40 ppm) were injected and detected at 250 °C. Initial oven temperature was 85 °C for 1 min and then increased to 190 °C at 2 °C/min. For both chiral identification and field assays, we used the (*E*)-nerolidol from Sigma-Aldrich, which was previously determined to be comprised of almost 100% (*3S,6E*)-nerolidol [21], and also confirmed in our GC analysis (Fig 3d).

### Field bioassays of synthetic flower volatiles

We conducted three independent field trials. In the first tests, we mimicked the *L. siceraria* flowers by using flower-shaped dummies made from white filter paper, which were impregnated with 100 mg of neat nerolidol (a mixture of isomers, Sigma-Aldrich). Control dummies had no compound. Dummies were attached to wood sticks (25 cm long), which were inserted into the ground, so the dummies were at 23 cm hight. Nerolidol-treated and control dummies were randomly distributed in three pairs, with each pair containing one dummy of one treatment. Dummies were spaced 3 m apart, and each pair of dummies was placed ∼ 5 m from the other. Bottle gourd flowers dummies were checked for visiting beetles from 06:00 p.m. to 09:00 p.m. Soon after landing, Cyclocephalini beetles were removed from dummies to avoid potential intraspecific attraction. This bioassay was carried out in the same *L. siceraria* groove in Piracicaba, SP, Brazil (see above) for 2 d (March 12-13, 2018), with dummies replacement made for each replicate.

The second field tests were conducted in the same area in Piracicaba and a pasture area owned by the Mato Grosso do Sul State University, Cassilândia, MS, Brazil (∼ 800 km from Piracicaba; 19°05’35.8” S, 51°48’52.5” W). We used custom-made, cross-vane intercept traps (black polystyrene panels: 18 × 24 cm), with 0.5-L plastic jars attached to the bottom of the trap. Both control and treatment traps were loaded with 200 ml of a solution of water and dish detergent to kill the captured Cyclocephalini beetles. Traps were hung from L-shaped PVC pipe frames (1.5 m long) that were supported by 1-m reinforcing steel bars hammed into the ground. The lures were pieces of filter paper (2 × 6 cm) loaded with 50 mg of neat nerolidol (a mixture of isomers). Control lures were filter papers with no compound. Lures were hung in the central opening slot of traps with a diaper pin 1-h prior to the sunset. Treated and control traps were randomly distributed. Seven even pairs of traps were deployed in Piracicaba and 10 pairs in Cassilândia. Traps were spaced 20 m apart, and pairs of traps were set ∼30 m from each other. Traps were serviced for captured beetles in the morning. For every replicate date, the lures were replaced, and the treatment traps were switched in position to avoid positional effects. Overall, the bioassay was repeated for 5 d in Piracicaba (October 17-22, 2018) and for 1 d in Cassilândia (October 1^st^, 2018).

The third bioassay was conducted in the same area in Cassilândia (October 1-2, 2019), which tested the following treatments: (i) 5 mg of neat (3*S*,6*E*)-nerolidol (*trans*-nerolidol, Sigma-Aldrich); (ii) 20 mg of neat nerolidol (a mixture of isomers), considering 5 mg per isomer; and (iii) control (filter paper with no compound). Treatments were assigned randomly to traps within five blocks. Evaluations were carried out every morning for two days. Lures were replaced 1-h before the sunset, and treatment traps were changed one position within blocks to control for positional effects.

Captured cyclocephalini specimens were sent to Dr. Paschoal Coelho Grossi (Federal Rural University of Pernambuco - UFRPE) for taxonomy. A voucher specimen of Cyclocephalini beetle, *C. paraguayensis*, was deposited in the collection of the Museum of Entomology (#6945), Department of Entomology and Acarology (University of São Paulo, Piracicaba, Brazil). Collection of beetles in the field were conducted under SISBIO permit #60104 and #67772 from the Brazilian Ministry of the Environment. This work was submitted to the Genetic Heritage Management Council by SisGen within project registration #AFC4FA2.

### Statistical Analysis

Data from experiments 1 and 2 were analyzed with the two-tailed Exact Binomial test to verify whether the ratio of beetles attracted to nerolidol and control differed significantly from the expected 1:1 ratio at 5% probability [22]. The test was performed from a spreadsheet provided by http://www.biostathandbook.com/exactgof.html (accessed January 17, 2019).

Differences between treatment means from experiment 3 were analyzed using the nonparametric Friedman’s Test (PROC FREQ, option CMH; SAS Institute, 2011), because data did not pass the assumptions of ANOVA [23]. Replicates were defined by block and collection dates. Replicates with no capture of *C. paraguayensis* in any trap were excluded from analyses. Once an overall significance was detected by the Friedman’s test, pairs of means were compared using the REGWQ multiple range test, which controls for type I experiment-wise error rate [24]. For all bioassays, the sex ratio of attracted adults of *C. paraguayensis* was compared to a nominal proportion of 0.5 with 95% Clopper-Pearson Exact Confidence Intervals [25].

## Acknowledgments

We thank Dr. Wolmar Trevisol (Federal University of Santa Maria, CAFW) for sharing *L. siceraria* seeds and Dr.Robert Hanus (Academy of Sciences of the Czech Republic) for a generous gift of enantiopure (*3R-6E*)-nerolidol. Special thanks to Lucila Wadt, Emiliana Romagnoli, Diego Silva, Fernando Madalon, Bruno Ribeiro, Hugo Rainho, and Francisco Prata for their invaluable help during the experiments, and Dr. Paschoal Grossi for identifying the beetle specimens.

